# Flexible Motion of T7 bacteriophage Tail Fibers Suggest a Dynamic Viral Infection Mechanism

**DOI:** 10.1101/2025.10.22.683886

**Authors:** Luca Elizabet Kosik, Miklós Cervenak, Dominik Sziklai, Andrea Balogh-Molnár, Negar Rahimi, Bence Fehér, Soma Yamamoto, Hiroki Konno, Noriyuki Kodera, Heinz Amenitsch, Hedvig Tordai, Levente Herényi, Miklós Kellermayer, Bálint Kiss

**Author notes:** Corresponding author: Balint Kiss.

## Abstract

Viruses are nanoscale infectious agents capable of specifically targeting and reprograming host cells. A unique group of viruses, bacteriophages, have regained popularity in research partly due to the rising number of multidrug-resistant bacterial infections. Phages could potentially replace antibiotics, but only if we understand every detail of their structure and infection cycle. T7 bacteriophages are a group of dsDNA viruses, which infect *E. coli* bacteria. T7 virions are comprised of an icosahedral protein shell which encapsulates the genomic DNA, and a tail-fiber complex which is primarily used for target recognition and DNA injection. The virus has six” L”-shaped, ∼40 nm long fibers (gp17 protein trimers) attached to the tail-tube, which are thought to be essential for initial host recognition and possibly surface exploration. Using high-speed atomic force microscopy (HS-AFM) and molecular dynamics (MD) simulations combined with small angle X-ray scattering (SAXS) we observed the molecular structure and movements of isolated tail fibers. Firstly, we have identified a hinge region within the fibers, which makes them highly flexible, allowing the bending of their distal region. Furthermore, we have observed the dynamic triple helical coiled coil structure of the proximal region, which would allow fiber rotation. These two points of flexibility allow a more efficient and highly dynamic host recognition and virus anchoring process. The observed flexibility might allow host surface exploration by walking. Such flexibility in the host recognition machinery may not be unique to T7 bacteriophages, getting us one step closer to understanding the intricate details of virus-host interactions.

## Introduction

The number of multidrug resistant bacterial infections are on the rise, posing an ever more pressing issue in our healthcare system ^1^. Instead of using conventional antibiotics, bacteriophages can provide a much needed and effective alternative, given that bacteriophages have evolved to infect bacteria with high specificity ^2, 3^. Phages can only be successfully modified and applied as an alternative therapy if we understand their infective cycle down to the smallest detail ^4^. Bulk experiments fail to unravel such details; only single particle approaches can shed some light on the individual steps of target recognition and infection ^2^. A workhorse of virus research, the T7 bacteriophage has long been of great interest, yet the exact process of their target recognition and infection mechanism remains unclear ^5^. T7 phages are formed by an icosahedral capsid, which encompasses the viral genome, and a tail fiber complex which is used for host recognition (*E. coli*) and genome delivery ^6^. Tail-fiber complexes consist of a tail-tube and six fibers ^7^. The tube itself is a complex of three rings formed by three different proteins (gp8, gp11and gp12), which extends from a vertex of the capsid ^8^. Gp8 is found inside the capsid, gp11 is serving as an adapter, connecting the inner gp8 ring to the outmost gp12 ring, which also serves as a gatekeeper of the viral DNA. Tail fibers are trimers of the gp17 protein, which connect to the tail-tube via their N-terminus between the dodecamer gp11 and hexamer gp12 rings. The fibers themselves have an overall “L” shaped structure. The two straight parts of the L-shape are called proximal and distal regions, according to their closeness to the tail-tube ^9^. Most of the distal region’s structure has been resolved by X-ray crystallography, whereas the proximal region’s structure remains largely unknown ^10^. Fibers are used for target recognition and anchoring the phage to the host cell lipopolysaccharides during DNA translocation (LPS) ^5, 11^. It is so far unknown whether it is the fibers or the tail tube that initially bind to the host cell.

Tail fibers were shown to be in two distinctly different conformations depending on the infection state of the bacteriophage ^5^. After new phage progeny are formed and released from the host cell via bacterial lysis, they start free diffusion in their surrounding medium ^12^. During this stage the fibers of free viruses were found to be in a capsid bound laid back orientation ^5, 13^. It is hypothesized that this conformation of the fibers is structurally more secure, as they are less likely to be accidentally broken off by random collisions with their surroundings. Following target recognition, phages extend their fibers towards the bacterium by a currently unknown mechanism. During infection, fibers are in an extended state, with a ∼90° internal bending angle between the proximal and distal region. This value, however, was calculated from averaged cryo electron microscope images, potentially obscuring phage-to-phage variability ^5^. We have recently found that phages can reversibly bind to the host cell surface, while we have also shown proof of host cell surface exploration ^13^. Such exploration was also observed in case of other virus species, like coronaviruses ^14^. The exact mechanism of this process is so far unknown, our recent findings have suggested exploration either by rolling or by walking, however we did not have the means to directly support one or the other mechanism.

In the current study we provide a detailed insight into the structure of the T7 bacteriophage. Prior observations have shown the presence of an elbow-like bend in the tail fiber. Here we demonstrate that fibers display a highly dynamic conformational flexibility along the elbow region on the second timescale. Furthermore, based on our results we describe the possible function of the coiled coil proximal region in fiber extension. Finally, our findings also provide details which support the possibility of viral host cell surface exploration by walking.

## Results and Discussion

### Virus associated and isolated tail fibers have a similar structure

Viral host cell recognition and infection processes have always been an extensively studied topic. Recent advances in AFM methodology can now offer insight on the level of individual nanoscopic objects, such as viruses in their native state ^15^. To get a general understanding of T7 morphology, whole T7 bacteriophages were scanned by atomic force microscopy (AFM) (Fig. 1a). Phages appear as icosahedral, 60 nm high particles, formed by capsomeres. Depending on the orientation of the phages on the substrate surface, one, two or in rare cases three ‘L’ shaped tail fibers of a single phage are visible. Fibers are clearly visible on the surface of capsids (Fig. 1a) and can always be found in a clockwise orientation. Proximal and distal regions are clearly distinguishable, enclosing a bending angle close to 90°. The average and standard deviation of fiber bending angles were calculated to be 94° ± 11° (AVG ± SD, n=45). Contrary to popular depiction of bacteriophages having extended fibers, they were always found in a secure, capsid bound orientation. Even though extended tail fibers were never visualized, we know that they play a crucial role in target recognition and in stabilizing the virus during DNA translocation ^16^. Fibers mostly being in a laid back, capsid attached orientation in case of freely diffusing bacteriophages was also noticed by *Hu et al* using cryo electron microscopy ^*5*^. A schematic representation helps the understanding of the structure of T7 phages and its laid back, virus attached tail fibers (Fig. 1d).

**Fig. 1.**
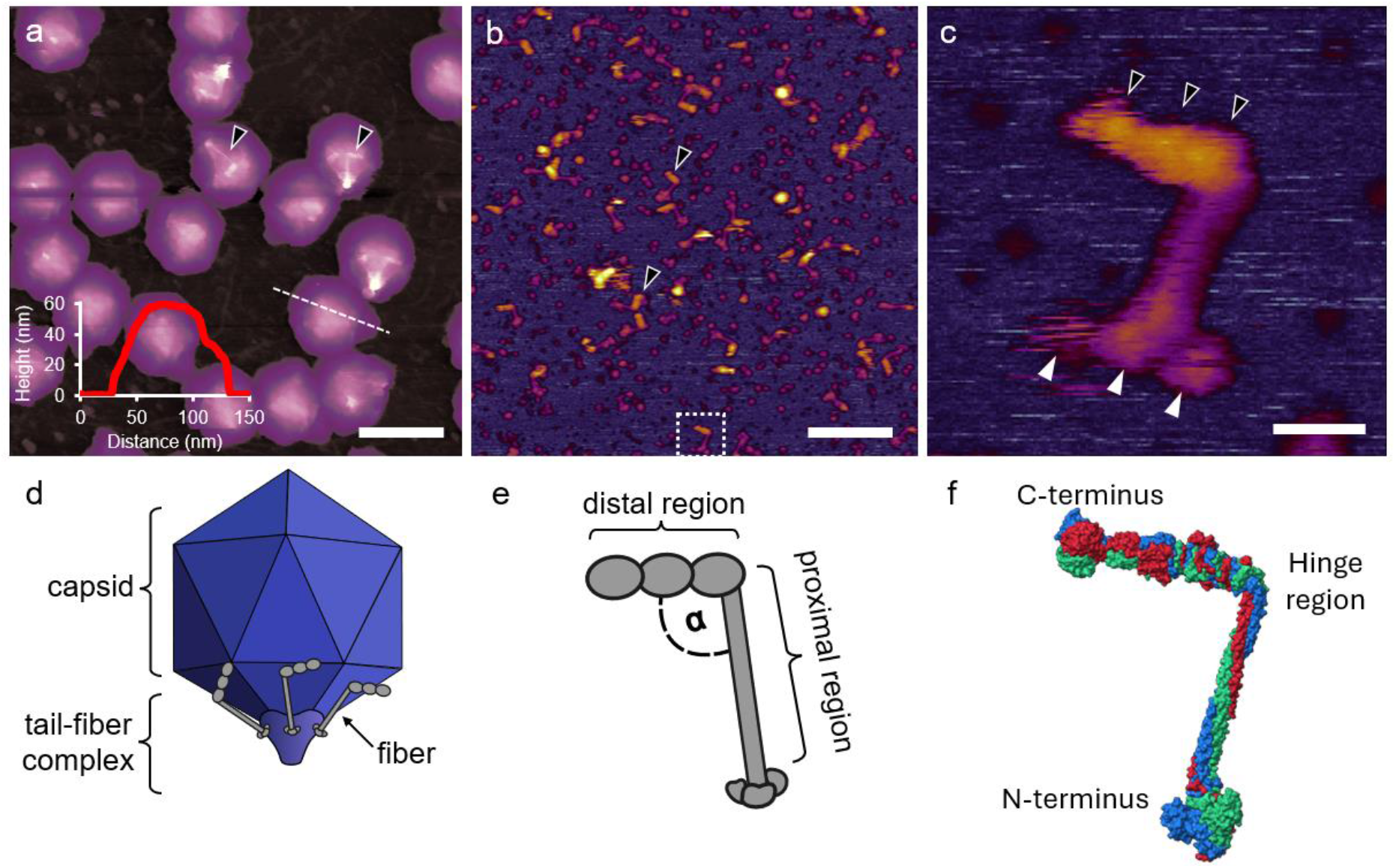
Tail fibers have an ‘L’ shaped conformation. **a** Height-contrast AFM image of complete T7 bacteriophages. Phages are shown in purple color; tail fibers are shown in white. Black arrowheads point at virus-attached tail fibers. Inset shows the height section profile of a single phage along the white dashed line. The scale bar is 100 nm. **b** Height-contrast AFM image of purified, substrate-adsorbed tail fibers. Black arrowheads show tail fibers. White dashed square shows a fiber of interest, enlarged in panel c. The scale bar is 100 nm. **c** Image of a single tail fiber in the more frequent, hence more probable, counterclockwise orientation. Black arrowheads show the three distal globular domains. White arrowheads show the three proximal globular domains. The scale bar is 10 nm. **d** Schematic representation of a T7 bacteriophage with its tail fibers in laid back orientation. **e** Schematic representation of a tail fiber, showing its different regions. **f** Final state of an equilibrium MD simulation of the viral fiber. Red, blue and green colors show individual monomers of gp17. The intermediate structures of the MD simulation are shown in Supplementary Movie 1.

Scanning fibers on the surface of bacteriophages is quite challenging, for this reason we have decided to recombinantly express tail-fibers. Imaging purified tail fibers on substrate surfaces has also revealed ‘L’ shaped structures (Fig. 1b). In the absence of the tail tube (gp-8-11-12) fibers do not form complexes with one another. Contrasted with virus-attached fibers, isolated ones are regularly found in a counterclockwise orientation (93% of analyzed fibers, n=208). Even though AlphaFold predictions show a largely homogeneous positive surface charge (Supp. Fig. 1), this change in orientation could arise due to an asymmetrical charge distribution on the two sides of the fibers. Alternatively, because the APTES coated surface and the viral capsid have different electrostatic properties. APTES being positively charged, whereas the surface of the capsid is negatively charged ^17^. The latter hypothesis seems more plausible, explaining the reason of capsid attached fibers. The length of the proximal region of the recombinant tail fibers was measured to be 24.0 ± 2.8 nm, while the distal region’s length was 15.4 ± 2.6 nm (AVG ± SD, n=208), which agrees well with already reported measurements ^9^. Proximal regions are thinner (∼4 nm), while distal regions are somewhat thicker (∼5 nm). A schematic representation displays the structure of the fiber showcasing its different regions (Fig. 1e). The proximal end of the fiber often appears as three separate globules, which are the N-termini of the three gp17 monomers, normally connected to the tail tube (gp8-11-12 complex) of the phage (Fig. 1c). These three globules are either displaced, or they form a single larger globule. The proximal region appears as a straight rod-like region, even though modeling has already suggested that it might contain a triple helical structure formed by the three monomers of gp17 (Fig. 1f). The thicker distal region of the fiber is formed by three sequential globular domains, which can also be seen in the acquired images. It is already apparent from AFM images that the angle enclosed by the proximal and distal regions can vary from fiber to fiber, ranging from 63° to 153° (94° ± 7°, AVG ± SD, n=208). To examine this broad angle distribution, we have performed AlphaFold modeling, molecular dynamics (MD) simulation and small angle X-ray scattering (SAXS) measurements.

### Fibers have a flexible hinge region

The most accurate AlphaFold structural prediction exhibits a straight conformation of the tail fiber (Fig. 2a). Predicted local distance difference test (pLDDT) scoring shows high confidence in the distal, as well as in the proximal region. However, their connecting region has a much lower confidence score, highlighting a potentially dynamic region with a less reliable structural prediction. This is in agreement with AFM images showing fibers with a ∼90° bend within their structure. The straight structure was used as an initial state for equilibrium all-atom molecular dynamics (MD) simulations, which all stabilized in an ‘L’ shaped structure after ∼300 ns of simulation (Fig. 2b). Analysis of the simulation trajectories identified two flexible regions. One of them is close to the N-terminus of the trimer, we call this the shoulder joint region. The second flexible region, which is of main interest for us, is the bent segment separating the proximal and the distal region, from here on called the hinge region. During the simulation most of the conformational changes are taking place in this region, which leads to the bending of the fiber (Fig. 2c). This bent shape seems to be the most stable conformation, as also confirmed by AFM scans, as well as longer simulations (>300 ns). Fiber bending angles from simulated structures were calculated to be 114° ± 21° (AVG ± SD) (Fig. 2e). Furthermore, torsion of the proximal region was also monitored, showing minimal standard deviation (11°) throughout the simulations.

**Fig. 2.**
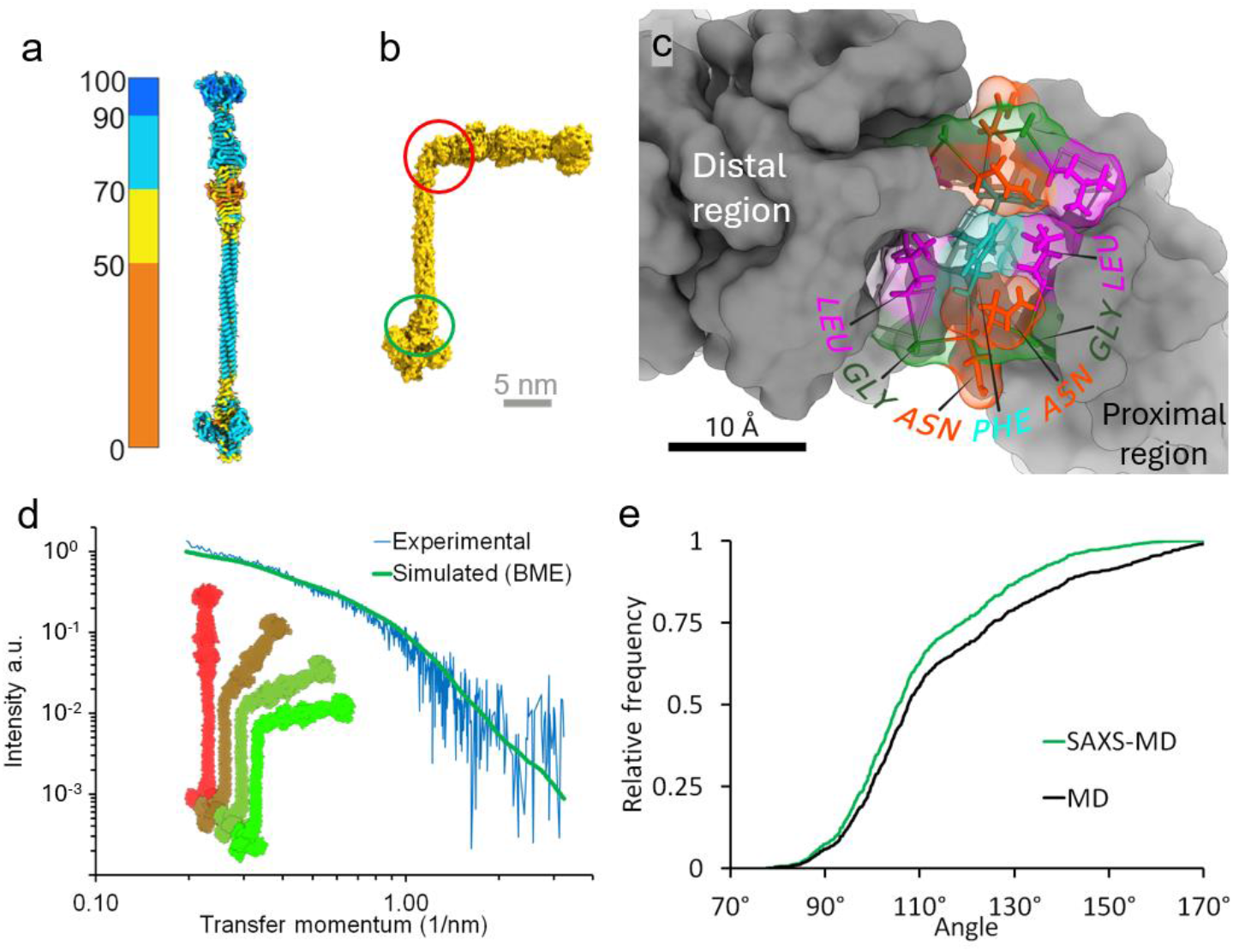
SAXS and MD simulations confirm the presence of a hinge region within the fiber. **a** AlphaFold3-predicted structure of a gp17 trimer, color bar indicates pLDDT scoring, blue shows high confidence, orange colors show low confidence regions. **b** Resulting structure of an equilibrium MD simulation, red circle highlights the hinge region of the structure shown in the c panel. Green circle highlights the shoulder joint region. **c** Structure of the hinge region (all three protein strands are shown) with the constituent amino acids of one of the protein’s, labeled with their three-letter codes. **d** X-ray scattering profile shown in a double logarithmic plot, displaying experimental and BME guided fitting curves. Inset shows different fiber conformations with red-to-green color scale, red being a less likely, green being the most likely conformation. **e** Cumulative distribution of the bending angle enclosed by the proximal and the distal regions of the fiber derived from MD simulation (black) and SAXS weighted structural fitting (green).

To experimentally calculate the bending angle distribution of fibers in a non-immobilized state we performed small-angle X-ray scattering (SAXS). SAXS is a powerful technique for characterizing the conformational ensembles of dynamic proteins, particularly when combined with all-atom or coarse-grained MD simulations. The experimental SAXS profile is shown in Fig. 2d, blue line. The scattering curve exhibits a slight upturn in the low-q region, suggesting the presence of a small fraction of aggregates. Primary data evaluation followed the guidelines of Jacques *et al*., and the resulting parameters are summarized in the Supplementary ^18^. Notably, the molar mass derived from the forward scattering (182 kDa) agrees well with the theoretical value of the gp17 trimer calculated from the amino acid sequence (189 kDa), providing a robust basis for further analysis. The most probable conformations (out of 750 conformations) were identified using the Bayesian maximum entropy (BME) approach, which assigned appropriate weights to each frame. Based on these representative conformations and their associated probabilities, the cumulative angle distribution was computed (109° ± 13°, AVG ± SD) (Fig. 2e). The MD-derived distribution closely matches the SAXS-refined distribution, although the experimental data indicates smaller bending angles. We can either contribute this to surface binding effects, or it might also indicate that fibers in their extended state, which can be more accurately modeled by free fibers, have a greater and more flexible bending angle.

### Proximal regions are formed by a dynamic coiled coil structure

AlphaFold predictions and MD simulations both point at the presence of a coiled coil region within the proximal region of the fiber. To our knowledge the structure of this region has not been resolved experimentally due to its presumed flexibility. We show for the first time that using high resolution AFM the dynamically changing proximal region can be resolved (Fig. 3). The seemingly rod-like section of the proximal region is in fact formed by a triple helical region with a pitch that can vary from fiber to fiber. The three monomers of gp17 can either form a tightly wound helical region (Fig. 3a) or they can sometimes be found in an almost straight conformation (Fig. 3b), where the three strands run parallel to one another. The whole population of fibers consists of conformational states between these two extremities (Fig. 3c-f). In case of virus connected tail fibers the loosening of the proximal region coiled coil would allow the rotation of the distal region. This process might be necessary for fiber extension, as well as host cell surface explorations.

**Fig. 3.**
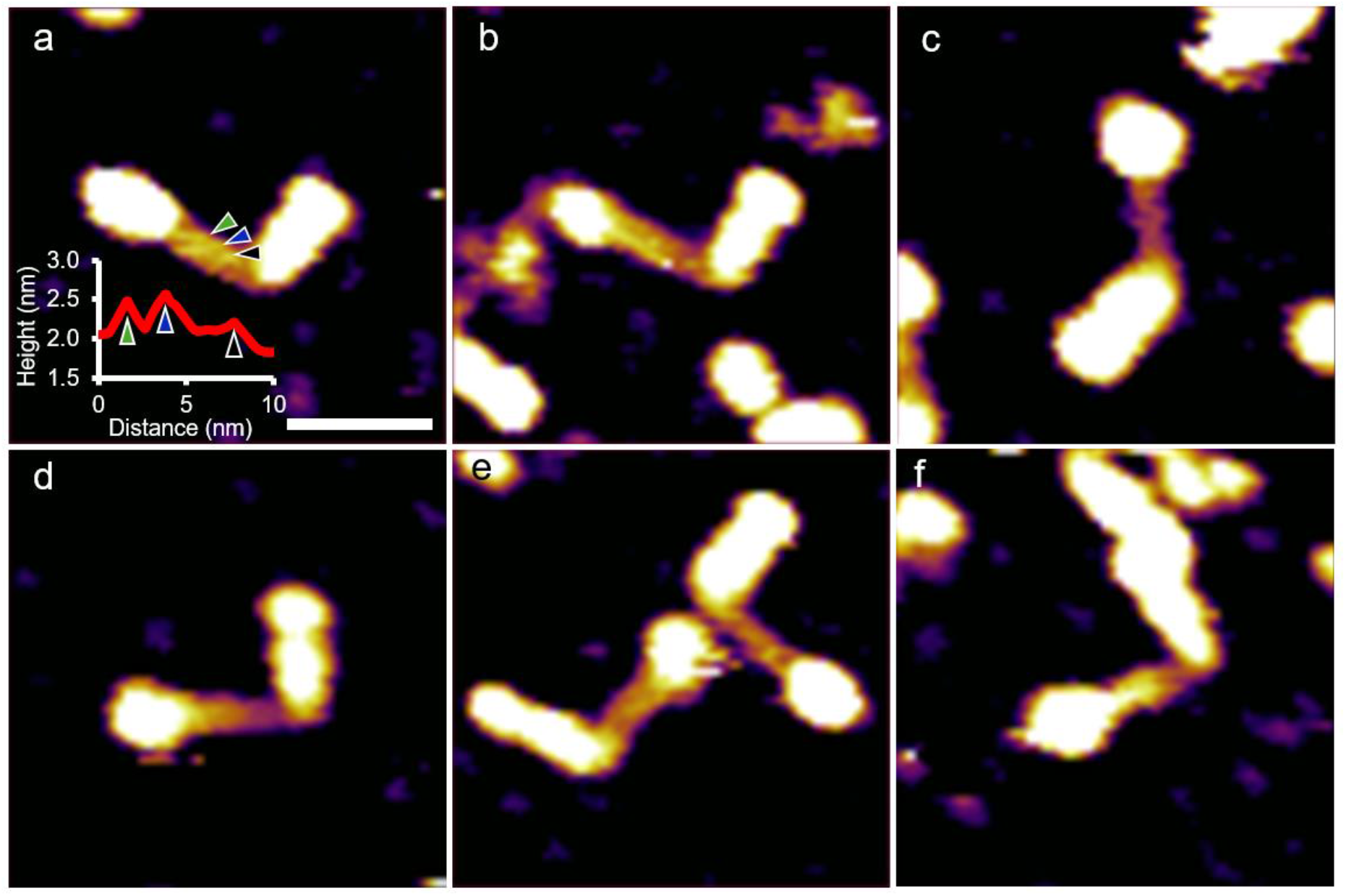
Fiber proximal region is a dynamic coiled coil. High resolution AFM height-contrast images of T7 fibers. **a** Fiber with a tightly wound triple helical proximal region. Inset shows height-section profile along the central line of the coiled coil proximal region. Colored arrowheads show the corresponding locations of the proximal region protein strands. **b** Fiber with a coiled coil region. **c-f** Fibers showcasing different degrees of pitch. Scalebar is 20 nm.

### Fiber bending visualized by high-speed AFM

Experimental results of the above-mentioned methods can only provide us with snapshots of the viral fiber’s conformation. To further investigate the hinge region’s flexibility and to gain insights into the movements of individual fibers, high-speed AFM (HS-AFM) experiments were conducted. Individual fibers were immobilized on substrate surfaces (see Methods section), then imaged at video rate scanning speeds (Fig. 4, Supplementary Movie 2). The overall structure of fibers appears similar to those observed by conventional AFM. At high framerates of 10 fps the movement of the distal arms was visualized (Fig. 4a). Initially, the fiber shown here is found with a bending angle close to 90° (Fig. 4a/1), which then transitions into a straighter conformation (Fig. 4a/2), followed by a quick transition back to a right-angle conformation. Fiber bending angles can change rapidly, within less than a tenth of a second from the more frequent close to right angle conformation to a straighter state (Fig. 4c). Bending angles of multiple fibers (n=11) were calculated frame-by-frame from HS-AFM videos, out of which the ones with longer observation times were selected (t > 20 s). Bending angle data was then pooled together, the cumulative distribution of which was visualized (97° ± 13°, AVG ± SD) (Fig. 4d). As discussed before, bending angle data was also collected from conventional AFM images of purified fibers, as well as from the fibers of whole bacteriophages. The distributions appear highly similar in all three AFM based measurements; angles vary between 70° to 180°. Interestingly, when compared to the MD and SAXS-MD results, both methods suggest a greater proportion of larger angles compared to AFM based angle calculations. We attribute this difference to the partial surface attachment of fibers during AFM imaging, which may restrict conformational flexibility. Nevertheless, the overall trends are highly consistent across different approaches.

**Fig. 4.**
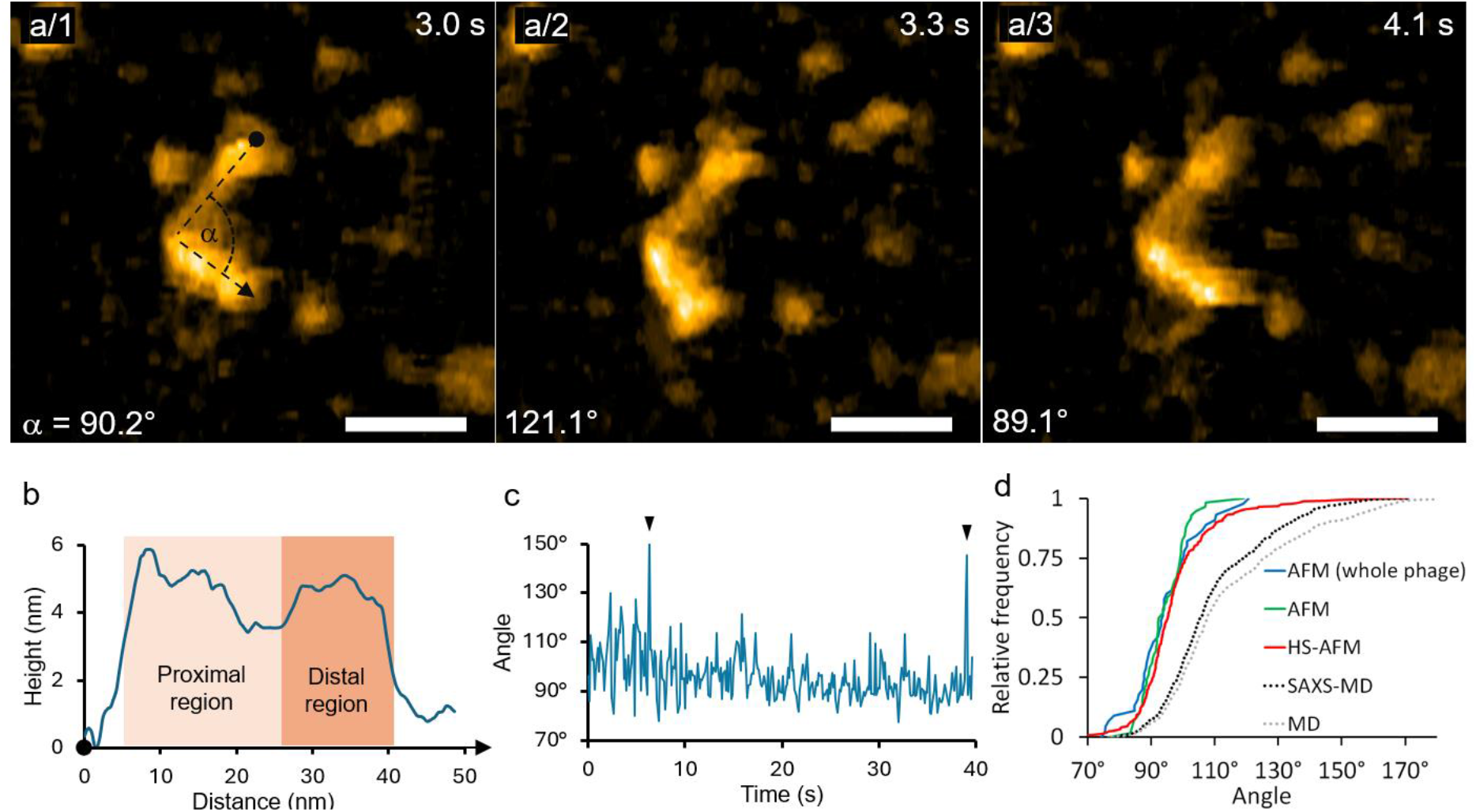
T7 tail fibers contain a flexible hinge region. **a** Height-contrast AFM image series of the same tail fiber (a/1 to a/3). Orange coloring shows the ‘L’ shaped fiber. Angles measured between the proximal and distal regions are shown in the bottom left (α), time stamps are shown in the top right. The scale bar is 20 nm in all three images. The related HS-AFM video can be seen in Supplementary Movie 2. **b** Height section profile of the fiber shown in a/1 along the black dashed line. Point and arrowhead show the direction of the measured height profile. **c** Angles between the proximal and distal regions of the displayed fiber measured over time. Black arrowheads show extreme cases of fiber bending. **d** Cumulative distribution of bending angles measured between the proximal and distal regions by multiple methods.

In rare cases fibers can even change into a straight conformation within less than 0.1 s, then quickly transition back to ∼90° bending angle in the same amount of time (Fig. 5, Supplementary Movie 3). Such rapid and extreme conformational changes were scarce. We would like to note that this quick conformational change closely resembles the MD simulations, although on a different timescale.

**Fig. 5.**
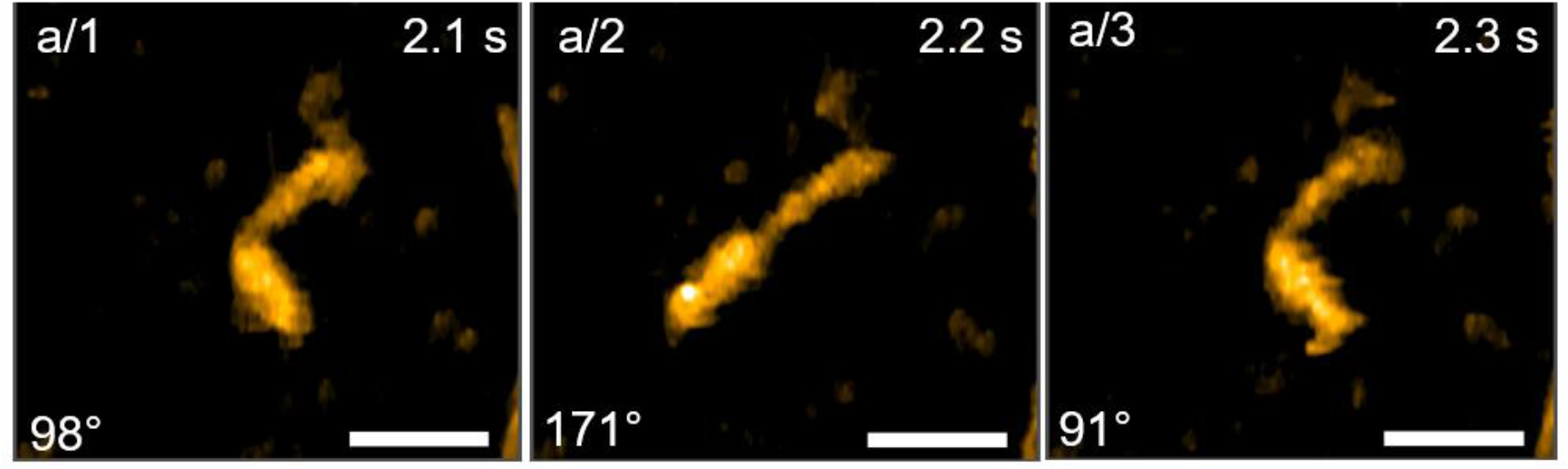
Tail fibers can occasionally straighten. **a** Height-contrast AFM image series of the same tail fiber (a/1 to a/3). Orange coloring shows the ‘L’ shaped fiber. Angles measured between the proximal and distal regions are shown in the bottom left (α), time stamps are shown in the top right. The scale bar is 20 nm in all three images. The related HS-AFM video can be seen in Supplementary Movie 3.

We have already recorded the movement of T7 bacteriophages on the surface of bacteria, confirming the process of viral host surface exploration ^13^. Our previous results, however, could only hint at the presence of such a process. The nanomechanical process of host cell exploration can be hypothesized by our current findings. The identification and characterization of the hinge region within the fiber would allow the conformational freedom necessary for viral exploration by “walking” (Fig. 7). Bending and straightening of the fibers might also drive the walking process. Bacteriophages may be able to explore an even larger region than the HS-AFM measured bending angles would suggest, since these measurements were conducted on partly immobilized fibers (resembling the capsid attached state). SAXS-MD shows a wider angular distribution compared to bending angles measured by AFM, and these fibers resemble the structure of extended fibers more closely due to their free diffusion state (Fig. 4d). Meaning, that during host cell surface exploration higher fiber flexibility could allow the exploration of a greater surface area. MD simulations also revealed a potential second hinge region within the N-terminal end of the fiber; however, this must be further explored, as the evaluation of HS-AFM images of this region are unreliable. It must be noted though, that walking would be further enhanced by the presence of a second hinge region at the base of the fibers.

### Fiber proximal region can uncoil over time

Conventional AFM images have already hinted at the possible dynamic rearrangement of the three globular domains at the end of the proximal region; as well as of the dynamic rearrangement of the proximal region’s coiled coil. Yet this method once again lacked the required time resolution to visualize such rearrangements. Using HS-AFM the rapid rearrangement of the N-terminal globular domains and the unwinding of the proximal coiled coil region was captured (Fig. 6a, Supplementary Movie 4). Initially, the presented fiber’s N-terminal end of the proximal region appears as a single 6 nm diameter globule (Fig. 6a/1). Over the span of a few seconds, the N-terminal domains dissociate, and the three smaller globular domains of the monomers become visible (Fig. 6a/2). These globules are most likely the ends of the gp17 monomers, which are normally bound to the tail-tube complex (gp8-11-12). The three globules can be seen wiggling around, presumably rotating around each other, leading to the partial unfolding of the triple helical coiled coil proximal region (Fig. 6a/3). While this region can transiently refold, it completely unfolds over time (Fig. 6a/4). Unfolding terminates at the hinge region; hence the distal region’s structural integrity appears to be unaffected. Kymograph and schematic illustrations help the understanding of the unfolding of the proximal region (Fig. 6bc). It must be noted that in its native state the N-terminal end of the fiber is attached to the tail-tube complex, between the gp11 dodecamer and the gp12 hexamer regions ^8^. In its isolated state the fiber’s N-terminal region may be quite unstable without binding to the tube.

**Fig. 6.**
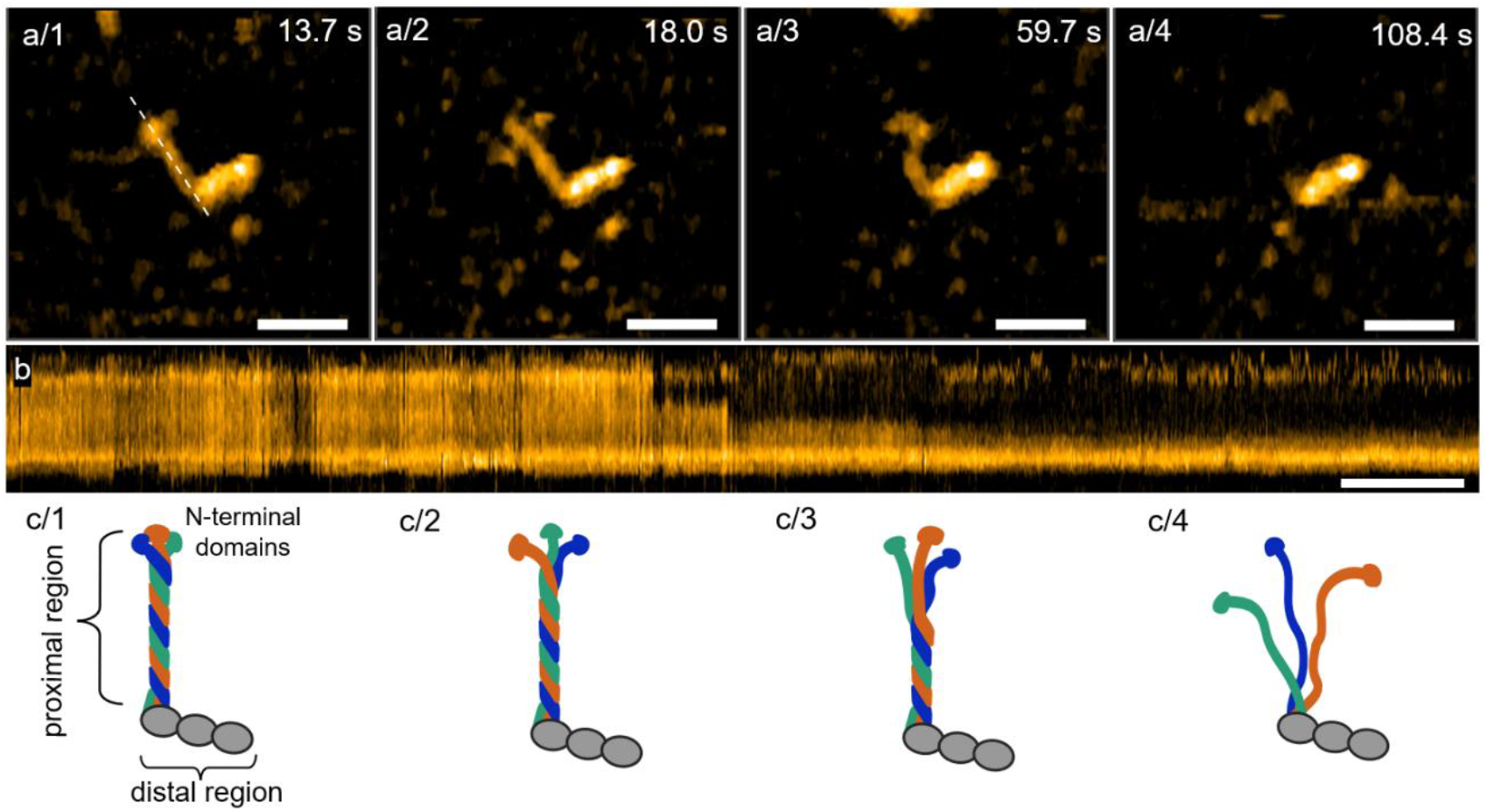
Proximal region unfolding over time. **a** AFM height-contrast image series of the same tail fiber, showing the unfolding of the proximal domain. a/1 has a complete triple helical proximal region, while in a/4 it is completely unfolded. White dashed line in a/1 shows the location of the kymograph displayed in b. The scalebar is 20 nm in all four images. **b** Kymograph of the proximal region’s unfolding, the scale bar is 10 seconds in the horizontal direction. **c** Schematic image series representing the states of proximal region unfolding shown in the panels (a/1 to a/4) above them, respectively. The related HS-AFM video can be seen in Supplementary Movie 4.

The proximal region of the fiber according to our modeling predictions and high-resolution AFM imaging is a coiled coil structure. Furthermore, using HS-AFM we have imaged the unfolding of the proximal region, which is most likely due to the unwinding of the coiled coil structure (Fig. 6). Such unwinding is only possible if the N-terminal globular regions can freely rotate. Thus, we propose a model, where the circular rotation of these regions unwinds the structure (Fig. 6c). Unfolding is proceeding rather slowly, complete proximal region unfolding happens over a span of at least a minute. The distal region however has a different, more stable quaternary structure, since the unwinding always stopped at the hinge region. We do not exclude the possibility of complete proximal region refolding; however, it did not occur in any of the recorded videos. The process of at least partial unwinding of the proximal region may play an important role in fiber extension as well as in surface exploration. For the fibers to be released from the capsid bound orientation and come down to an extended state a conformational change of the tail fiber complex must be initiated. It is also important that the fibers need to rotate by ∼90° to touch the bacterial surface (Fig. 7). We propose a model where the proximal coiled coil might act as a torsional spring, capable of storing mechanical energy. We can only speculate about how this torsional stress was created to begin with during phage production. It is possible that similar to mechanisms seen in contractile tail phages, the whole tail-tube might reorganize and rotate as a final step of virus formation ^19^. This process might involve the reorganization of the gp12 ring. Since the base of fibers (N-termini) are connected between the gp11 dodecamer and gp12 hexamers rotation of the gp12 ring region could force the fibers to transition to their capsid bound state. Similarly, upon receptor recognition the gp12 ring might rotate again, but in the other direction forcing the fibers into their extended state. Alternatively, fibers could be extended one by one, as a result of coming in contact with the bacterial surface.

**Fig. 7.**
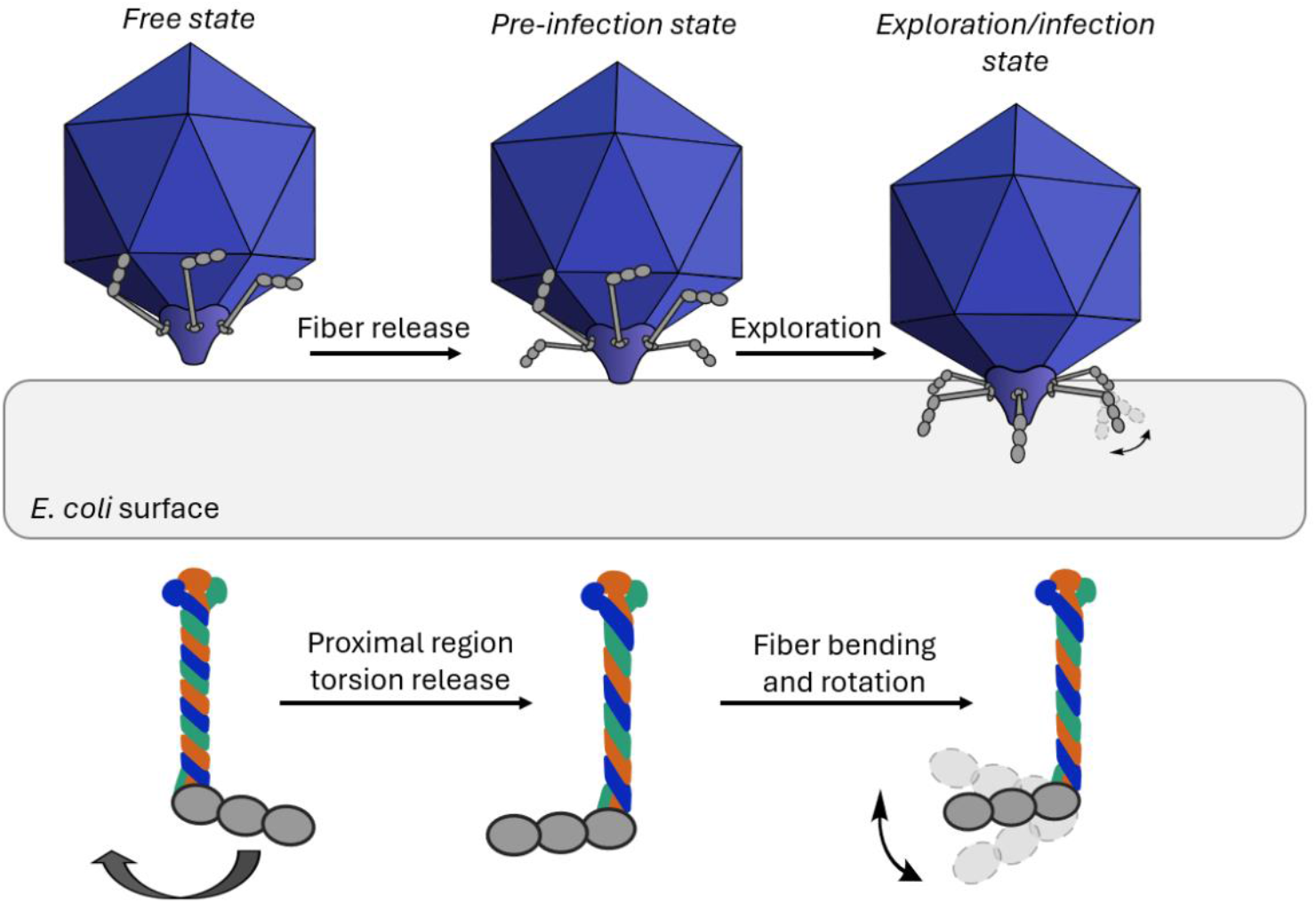
Viral fibers extended by rotation along the proximal region, walking and receptor searching enabled by the presence of a hinge region.

Another probable explanation is that when the fibers come into close proximity to the bacterial surface its electrostatic screening effect and possibly stronger negative charge (compared to the capsid) might trigger their extension on the side that is touching the bacterium. This process could potentially be followed by viral surface exploration by walking, for which both fiber bending at the hinge and rotation along the proximal coiled coil is crucial (Fig. 7). This process would also involve reversible binding of the fibers to the LPS layer. Finally, for all six fibers to be extended, flexibility is once again crucial, as fibers on opposite sides of the tail can only come into contact with the host surface if the already attached fibers extension allows it. Our experiments shed some light on fiber release as well as viral surface exploration, nevertheless, experiments involving the whole tail fiber complex could provide even more specific details about the viral host cell recognition process.

### Conclusion and outlook

Viral host cell recognition, surface exploration, and infection were always of great interest; however, certain steps of the process are not yet clear. Our results hint at the possible mechanisms of fiber extension by a torsionally loaded spring mechanism. Furthermore, we propose a model, whereby T7 bacteriophages explore the host cell surface by a “walking” mechanism enabled by the fiber’s internal flexibility. These processes may not be exclusive to the T7 phage, many types of bacteriophages and human viruses might utilize such mechanisms during host recognition and target infection.

## Materials and Methods

Purified T7 bacteriophages were a kind gift from the group of Gabriella Csík ^20^. One Shot™ BL21(DE3) pLysS Chemically Competent *E. coli* (Thermo Scientific) was transfected with gp17 coding plasmid, which was a kind gift from Mark van Raaij. Lysogeny broth (LB), glutaraldehyde grade I (GA) solution, poly-L-lysine (PLL) solution, 3-Aminopropyl triethoxysilane (APTES), IPTG, imidazole, and basic chemicals were purchased from Merck. HisPur Ni-NTA Resin was purchased from ThermoFisher Scientific. Nitrogen 5.0 gas was from Linde Gáz Magyarország Zrt (Budapest, Hungary). Water was purified with a Milli-Q Integral 3 Water Production Unit (Merck Millipore, Billerica, MA). Round mica sheets were from Ted Pella, Inc. (Redding, CA).

### Expression and isolation of tail fibers

Preparations of tail fibers were done according to the work of *Garcia-Doval et al* ^21^. Briefly, cultures of BL21 *E. coli* were transfected with PET30(+) plasmid encoding the T7 tail fiber (gp17) protein. Transfected bacteria were grown overnight in LB growth medium containing kanamycin (50 µg/ml) and chloramphenicol (34 µg/ml). On the following day 4 x 250 ml growth media were inoculated with 4 x 2 ml overnight culture and grown while shaking at 37°C to OD600 ∼ 0.6. Expression was induced with 100 µM IPTG, cultures were then grown for 16 hours at 16 °C while shaking. Cells were collected by spinning them at 5000 g for 10 min. If necessary, pellets were stored at -80°C. Pellets were resuspended in 30 ml lysis buffer (50 mM TRIS, 4 % (v/v) glycerol, 150 mM NaCl, pH 8, supplemented with a protease inhibitor cocktail (Pierce, A32961)). Resuspended cells were lysed by sonication. The lysate was cleared by centrifugation at 38,000 g for 40 min at 4 °C. The supernatant was loaded onto a Ni-NTA column, after washing with a buffer containing 50 mM imidazole, the His-tag labeled fibers were eluted by a 500 mM imidazole buffer. 500 µl fractions were collected and flash frozen in liquid nitrogen, then stored at -80°C. Proteins were also found to be stable for at least three weeks at 4°C. Samples were further cleaned and concentrated using 100kDa Amicon Ultra centrifugal filters (Merck). Protein concentration of ∼ 2 mg/ml could be achieved by combining multiple fractions. The presence of tail fibers (gp17 trimers) was confirmed by SDS-PAGE.

### AFM experiments

#### Imaging of whole T7 bacteriophages

Surfaces for Atomic Force Microscopy (AFM) imaging were prepeared by dropping 100 µl of PLL onto freshly cleaved mica, followed by 10 min incubation, then washing with deionized water and drying in nitrogen stream. Then 100 µl of 25% (w/w) GA was dropped on top of the PLL coated surface, incubated for 20 min, followed once again by washing and drying. 100 µl of 100-200x diluted T7 bacteriophages (∼10^11^ PFU/ml) were dropped onto the PLL-GA coated mica and incubated for 30 min. Unbound phages were then washed away by repeatedly pipetting and removing 100 µl of PBS. All the above steps were performed at room temperature. Imaging was performed in PBS (137 mM NaCl, 2.7 mM KCl, 8 mM Na_2_HPO_4_, 1.5 mM KH_2_PO_4_) (pH 7.2) with an Oxford Instruments, Cypher ES scanner, using an Olympus BLAC-40TS cantilevers (typical spring constant of 90 pN/nm) in non-contact mode, with scanning speeds of ∼1 µm/s. Imaging was done at 25 °C.

#### Imaging tail fibers

AFM surfaces were prepared by dropping 100 µl of 10,000 x diluted APTES onto freshly cleaved mica, followed by washing with deionized water and drying in a nitrogen stream. AFM experiments were conducted by dropping 100 µl of 1,000-10,000 x diluted fiber samples (∼ 1-10 µg/ml) onto the APTES coated mica. Samples were incubated at room temperature for 5 min, then washed 5 x with PBS. Imaging was performed as described above.

#### High resolution imaging of tail fibers

APTES coated substrate surfaces were prepared as described above. 100 µl of 25% (w/w) GA was dropped on top of the APTES coated surface and incubated for 20 minutes, followed by washing with deionized water, then drying in N_2_ stream. Fibers were incubated on this surface and washed as described above. Imaging was carried out in PBS, using a Bruker PEAKFORCE-HIRS-F-B cantilever (nominal tip radius ∼1 nm) in non-contact mode utilizing a Cypher ES scanner. The sample holding area was flooded with ∼3 ml PBS, then closed and setup was left to equilibrate for 1 hour. (20°C).

### HS-AFM experiments

#### Imaging tail fibers

High-speed AFM (HS-AFM) substrate surfaces were prepared in a similar manner to conventional AFM experiments; however, imaging was performed in 3 x diluted PBS to optimize the degree of sample to surface binding and a small mica disk was mounted on top of a small glass stage which was directly glued onto the Z-piezo scanner. The sample stage was immersed in a cell containing around 50 μl of 3 x diluted PBS. HS-AFM imaging was performed in non-contact mode using a laboratory-built apparatus (WPI-NanoLSI, Kanazawa, Japan) ^22^. Scanning was done using custom made Olympus AC10 cantilevers with resonance frequencies of ∼200-600 kHz in water; with a typical spring constant of 100 pN/nm. Cantilevers were oscillated with 6-7 nm amplitudes. Scanning was performed with tip velocities of ∼100 µm/s on an 80×80 nm region, thereby achieving ∼10 frames per second.

#### AFM data analysis

Images acquired with Cypher ES were post processed and analyzed by the AFM driving software (AR 16 running in Igor Pro 6, Wavemetrics). HS-AFM data was post processed and analyzed using UMEX viewer developed by NanoLSI. Fibers were detected and analyzed using a custom in-house algorithm. Resulting data was sorted, processed and finally plotted using Origin and Microsoft Excel.

#### SAXS experiments and data analysis

SAXS measurements were performed at the Austrian SAXS beamline of the Elettra synchrotron (Trieste, Italy) in transmission geometry ^23^. Measurements were conducted using the 8 keV X-ray energy branch of the monochromator. Using a Pilatus3 1 M two-dimensional position sensitive CMOS hybrid pixel detector (Dectris Ltd, Baden, Switzerland), the scattering patterns were recorded in the range of 0.1–5 nm^−1^ in terms of the scattering vector, q (defined as 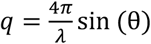, where 2θ is the scattering angle and λ is the X-ray wavelength), To be able to assess sample stability during the experiment, twenty exposures, each 10 s long, were made and repeated for each sample. The relevant aqueous medium was measured before and after the measurement series. The corrected scattering patterns were azimuthally averaged to yield one-dimensional scattering curves for each sample using SAXSDOG ^24^. The individual exposures were corrected for beam flux, geometric effects and for sample transmission. the µ-drop autosampler installed at the beamline ^24^. The background scattering curve was subtracted in the case of the measurements of fibers. The corrected scattering patterns were azimuthally averaged to yield one-dimensional scattering curves for each sample.

#### Molecular dynamics simulation and data analysis

Structural prediction of the gp17 (Uniprot ACC: H6BFI5) homotrimer protein complex was done using the AlphaFold Server ^25^. The multimer with the highest ranking-score (0.61) was used for subsequent simulation with a disordered fraction of 0.26, pTM of 0.5 and ipTM of 0.48.

All-atom molecular dynamics (MD) simulations were performed using GROMACS 2023.2 with the CHARMM27 force field ^26^. The initial protein structure was obtained from AlphaFold predictions and placed in a rectangular simulation box with edge lengths of 30, 35, and 20 nm. The system was solvated using the TIP3P water model. Sodium (Na^+^) and chloride (Cl^−^) ions were added to neutralize the net charge. Energy minimization was carried out using the steepest descent algorithm for 5,000 steps.

Equilibration was performed for 500 ps using a 2 fs integration time step. Long-range electrostatic interactions were computed using the reaction-field method with a cutoff distance of 1.2 nm and van der Waals interactions were treated with the same cutoff ^27^. Temperature was maintained at 310.15 K using the velocity-rescaling thermostat with a coupling time of 1 ps and pressure was isotropically controlled using the C-rescale barostat with a 5 ps coupling constant ^28^. All covalent bonds involving hydrogen atoms were constrained using the LINCS algorithm^29^.

Three independent MD simulation runs of 500 ns each were carried out with initial velocities assigned from a Maxwell–Boltzmann distribution using different random seeds. 250 equally spaced frames per trajectory were extracted, yielding a total of 750 conformations.

Theoretical SAXS profiles for each MD frame were computed using CRYSOL 3 ^30^. To estimate the optimal constant background level and scaling factor, CRYSOL was first run in fitting mode using the experimental SAXS curve for each structure. From each resulting file, the optimal background and scaling factors were parsed automatically. A second round of CRYSOL was then run without the experimental data, but using the extracted background and scale values, ensuring the resulting intensity files represented unbiased theoretical predictions adjusted for contrast matching. This step was parallelized for several hundred frames using a Python script.

To reconcile the conformational ensemble with the experimental SAXS data, Bayesian Maximum Entropy (BME) reweighting was performed ^31^. For each frame, the CRYSOL-generated intensity profile was interpolated to match the experimental q-range. The BME algorithm was then applied to compute an optimal set of statistical weights w_i_ for the frames by minimizing the Kullback–Leibler divergence D_KL_ from a uniform prior subject to agreement with the SAXS data within a confidence parameter θ. The effective ensemble was defined as:

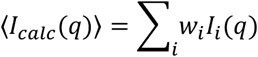

The weights were bootstrapped to estimate standard deviations.

Frames with nonzero weights were selected as representative structures contributing to the reweighted ensemble. To visualize structural convergence, the corresponding PDB files were loaded into PyMOL 2.

#### Measurement of angles and torsion in MD structures

Alpha C atoms of the multimer were extracted from the simulated protein structures using UCSF ChimeraX ^32^. Two linear fits were done on the extracted C atoms in Python using scikit-spatial (v6.8.1, 2019, Andrew Hynes): one fit on residues 160-250 and one fit on residues 280-558, thus excluding the regions where the structure bends (hinge region). Angles were measured between these two linear fits.

Torsion was calculated with the help of two planes. The first plane was defined at the end of the fiber by using the 480th serine residues of the three monomers. The second plane was defined with the help of the two linear fits (the two opposite ends and their intersections). Torsion was calculated as the angle between subsequent intersections of the two planes.

## Acknowledgements

First of all, we are thankful to Gabriella Csík, who provided for us the stock of T7 bacteriophages. The original plasmids encoding the tail fibers (gp17) were a kind gift from Mark van Raaij (Departamento de Estructura de Macromoléculas, Madrid, Spain), whom we would like to thank. We would also like to express our gratitude to Ana Cuervo, Tamás Hegedüs and Holger Flechsig for thought-provoking ideas and great talks. We are also grateful to Mónika Komárné Drabbant, Krisztina Lór and Zsófia Kovács for technical assistance. The authors acknowledge CERIC-ERIC consortium to access the TU Graz’s Austrian SAXS beamline@ELETTRA-Sincrotrone Trieste. This work was supported by the Hungarian Scientific Research Fund (K143321) and by HUN-REN-KUTT-2025 (02073), Biophysical Virology Research Group. This work was partly supported by Bio-SPM Collaborative Research program, World Premier International Research Center Initiative (WPI), MEXT, Japan and the Kanazawa University CHOZEN project. Balint Kiss was supported by the EKÖP-2025-328 university excellence scholarship program of the Hungarian Ministry for Culture and Innovation from the source of the National Research, Development and Innovation Fund. We acknowledge [KIFÜ] for awarding us access to resources based in Hungary at Komondor for CPU and GPU time and to HUN-REN Welcome Home and Foreign Researcher Recruitment Programme 2023.

## Author contributions

L.E.K., A. B-M., B.K. and H.T. prepared samples. L.E.K., M.C. and B.K. performed AFM experiments. S.Y., H.K., N.K., B.K. designed and performed HS-AFM experiments. D.Sz. and B.F. performed simulations. B.F. performed SAXS experiments and data analysis. L.E.K., B.K. and L.H. performed AFM data analysis. B.K. and M.K. conceived the project and wrote the manuscript. M.K. secured funding. All authors have read and approved the manuscript.

## References

(1) Jernigan, J. A.; Hatfield, K. M.; Wolford, H.; Nelson, R. E.; Olubajo, B.; Reddy, S. C.; McCarthy, N.; Paul, P.; McDonald, L. C.; Kallen, A. Multidrug-resistant bacterial infections in US hospitalized patients, 2012–2017. New England Journal of Medicine 2020, 382 (14), 1309–1319.

(2) Kiss, B.; Mudra, D.; Török, G.; Mártonfalvi, Z.; Csík, G.; Herényi, L.; Kellermayer, M. Singleparticle virology. Biophysical Reviews 2020, 12 (5), 1141–1154. DOI: 10.1007/s12551-020-00747-9.

(3) Mangenot, S.; Hochrein, M.; Rädler, J.; Letellier, L. Real-Time Imaging of DNA Ejection from Single Phage Particles. Current Biology 2005, 15 (5), 430–435. DOI: 10.1016/j.cub.2004.12.080.

(4) Viertel, T. M.; Ritter, K.; Horz, H.-P. Viruses versus bacteria—novel approaches to phage therapy as a tool against multidrug-resistant pathogens. Journal of Antimicrobial Chemotherapy 2014, 69 (9), 2326–2336. Lin, D. M.; Koskella, B.; Lin, H. C. Phage therapy: An alternative to antibiotics in the age of multi-drug resistance. World journal of gastrointestinal pharmacology and therapeutics 2017, 8 (3), 162.

(5) Hu, B.; Margolin, W.; Molineux, I. J.; Liu, J. The bacteriophage t7 virion undergoes extensive structural remodeling during infection. Science 2013, 1231887.

(6) Molineux, I. J. No syringes please, ejection of phage T7 DNA from the virion is enzyme driven. Molecular microbiology 2001, 40 (1), 1–8. Molineux, I. J.; Panja, D. Popping the cork: mechanisms of phage genome ejection. Nature Reviews Microbiology 2013, 11 (3), 194.

(7) Cuervo, A.; Fàbrega-Ferrer, M.; Machón, C.; Conesa, J. J.; Fernández, F. J.; Pérez-Luque, R.; Pérez-Ruiz, M.; Pous, J.; Vega, M. C.; Carrascosa, J. L.; et al. Structures of T7 bacteriophage portal and tail suggest a viral DNA retention and ejection mechanism. Nature Communications 2019, 10 (1), 3746. DOI: 10.1038/s41467-019-11705-9.

(8) Cuervo, A.; Pulido-Cid, M.; Chagoyen, M.; Arranz, R.; González-García, V. A.; Garcia-Doval, C.; Castón, J. R.; Valpuesta, J. M.; van Raaij, M. J.; Martín-Benito, J. Structural characterization of the bacteriophage T7 tail machinery. Journal of Biological Chemistry 2013, 288 (36), 26290–26299.

(9) Steven, A.; Trus, B.; Maizel, J.; Unser, M.; Parry, D.; Wall, J.; Hainfeld, J.; Studier, F. Molecular substructure of a viral receptor-recognition protein: The gp17 tail-fiber of bacteriophage T7. Journal of molecular biology 1988, 200 (2), 351–365.

(10) Garcia-Doval, C.; van Raaij, M. J. Crystallization of the C-terminal domain of the bacteriophage T7 fibre protein gp17. Structural Biology and Crystallization Communications 2012, 68 (2), 166–171.

(11) González-García, V. A.; Bocanegra, R.; Pulido-Cid, M.; Martín-Benito, J.; Cuervo, A.; Carrascosa, J. L. Characterization of the initial steps in the T7 DNA ejection process. Bacteriophage 2015, 5 (3), e1056904.

(12) Young, R. Phage lysis: Three steps, three choices, one outcome. Journal of Microbiology 2014, 52 (3), 243–258. DOI: 10.1007/s12275-014-4087-z.

(13) Kiss, B.; Kiss, L. A.; Lohinai, Z. D.; Mudra, D.; Tordai, H.; Herenyi, L.; Csík, G.; Kellermayer, M. Imaging the infection cycle of T7 at the single virion level. International Journal of Molecular Sciences 2022, 23 (19), 11252.

(14) Li, S.; Yang, H.; Tian, F.; Li, W.; Wang, H.; Shi, X.; Cui, Z.; Shan, Y. Unveiling the Dynamic Mechanism of SARS-CoV-2 Entry Host Cells at the Single-Particle Level. ACS Nano 2024, 18 (41), 27891–27904. DOI: 10.1021/acsnano.4c04212.

(15) de Pablo, P. J. Chapter Six - The application of atomic force microscopy for viruses and protein shells: Imaging and spectroscopy. In Advances in Virus Research, Rey, F. A. Ed.; Vol. 105; Academic Press, 2019; pp 161–187.

(16) Vörös, Z.; Csík, G.; Herényi, L.; Kellermayer, M. Temperature-dependent nanomechanics and topography of bacteriophage T7. Journal of virology 2018, 92 (20), e01236–01218.

(17) Gabashvili, I. S.; Khan, S. A.; Hayes, S. J.; Serwer, P. Polymorphism of bacteriophage T7. Journal of molecular biology 1997, 273 (3), 658–667.

(18) Jacques, D. A.; Trewhella, J. Small-angle scattering for structural biology—Expanding the frontier while avoiding the pitfalls. Protein Science 2010, 19 (4), 642–657. DOI: 10.1002/pro.351 (acccessed 2025/09/25).

(19) Leiman, P. G.; Shneider, M. M. Contractile tail machines of bacteriophages. Viral molecular machines 2011, 93–114.

(20) Gábor, F.; Szolnoki, J.; Tóth, K.; Fekete, A.; Maillard, P.; Csík, G. Photoinduced Inactivation of T7 Phage Sensitized by Symmetrically and Asymmetrically Substituted Tetraphenyl Porphyrin: Comparison of Efficiency and Mechanism of Action¶. Photochemistry and Photobiology 2001, 73 (3), 304–311.

(21) Garcia-Doval, C.; van Raaij, M. J. Structure of the receptor-binding carboxy-terminal domain of bacteriophage T7 tail fibers. Proceedings of the National Academy of Sciences 2012, 109 (24), 9390–9395. DOI: 10.1073/pnas.1119719109 (acccessed 2025/07/01).

(22) Ando, T.; Uchihashi, T.; Kodera, N. High-speed AFM and applications to biomolecular systems. Annual review of biophysics 2013, 42 (1), 393–414.

(23) Amenitsch, H.; Rappolt, M.; Kriechbaum, M.; Mio, H.; Laggner, P.; Bernstorff, S. First performance assessment of the small-angle X-ray scattering beamline at ELETTRA. Synchrotron Radiation 1998, 5 (3), 506–508.

(24) Haider, R.; Sartori, B.; Radeticchio, A.; Wolf, M.; Dal Zilio, S.; Marmiroli, B.; Amenitsch, H. µDrop: A system for high-throughput small-angle X-ray scattering measurements of microlitre samples. Applied Crystallography 2021, 54 (1), 132–141.

(25) Abramson, J.; Adler, J.; Dunger, J.; Evans, R.; Green, T.; Pritzel, A.; Ronneberger, O.; Willmore, L.; Ballard, A. J.; Bambrick, J. Accurate structure prediction of biomolecular interactions with AlphaFold 3. Nature 2024, 630 (8016), 493–500. Abramson, J.; Adler, J.; Dunger, J.; Evans, R.; Green, T.; Pritzel, A.; Ronneberger, O.; Willmore, L.; Ballard, A. J.; Bambrick, J.; et al. Accurate structure prediction of biomolecular interactions with AlphaFold 3. Nature 2024, 630 (8016), 493–500. DOI: 10.1038/s41586-024-07487-w.

(26) Pronk, S.; Páll, S.; Schulz, R.; Larsson, P.; Bjelkmar, P.; Apostolov, R.; Shirts, M. R.; Smith, J. C.; Kasson, P. M.; Van Der Spoel, D. GROMACS 4.5: a high-throughput and highly parallel open source molecular simulation toolkit. Bioinformatics 2013, 29 (7), 845–854.

(27) Tironi, I. G.; Sperb, R.; Smith, P. E.; van Gunsteren, W. F. A generalized reaction field method for molecular dynamics simulations. The Journal of chemical physics 1995, 102 (13), 5451–5459.

(28) Bussi, G.; Donadio, D.; Parrinello, M. Canonical sampling through velocity rescaling. The Journal of chemical physics 2007, 126 (1).

(29) Hess, R. A.; Bunick, D.; Lee, K.-H.; Bahr, J.; Taylor, J. A.; Korach, K. S.; Lubahn, D. B. A role for oestrogens in the male reproductive system. Nature 1997, 390 (6659), 509–512.

(30) Svergun, D.; Barberato, C.; Koch, M. H. CRYSOL–a program to evaluate X-ray solution scattering of biological macromolecules from atomic coordinates. Applied Crystallography 1995, 28 (6), 768–773.

(31) Bottaro, S.; Bengtsen, T.; Lindorff-Larsen, K. Integrating molecular simulation and experimental data: a Bayesian/maximum entropy reweighting approach. In Structural bioinformatics: methods and protocols, Springer, 2020; pp 219–240.

(32) Meng, E. C.; Goddard, T. D.; Pettersen, E. F.; Couch, G. S.; Pearson, Z. J.; Morris, J. H.; Ferrin, T. E. UCSF ChimeraX: Tools for structure building and analysis. Protein Science 2023, 32 (11), e4792.

