## Supplementary File for "Flexible Motion of T7 bacteriophage Tail Fibers Suggest a Dynamic Viral Infection Mechanism"

SAXS Data collection parameters

| | |
|---|---|
| Instrument | Austrian SAXS beamline@Elettra |
| Beam geometry | 0.5 x 1.5 mm transmission |
| Wavelength ( Angström) | 1.54 |
| q range (1/Angström) | $4.7 * 10^{-2} – 3.2$ |
| Exposure time (s) | 12x10 s |
| Concentration (mg / mL) | 0.5 |
| Temperature (K) | 303 |

Structural parameters

| | |
|---|---|
| I(0) (cm-1) [from P(r)] | $0.037 \pm 0.0023$ |
| Rg (nm) [from P(r)] | $4.72 \pm 0.04$ |
| I(0) (cm-1) [from Gunier fit] | $0.04 \pm 0.0017$ |
| Rg (nm) [from Gunier fit] | $4.70 \pm 0.04$ |
| Dmax (nm) | 14.43 |
| Porod volume estimate (nm3) | 150 |

Molecular-mass determination

| | |
|---|---|
| Calculated from I(0) (kDa) | 182 |
| Calculated from sequence (kDa) | 189 |

Software employed

| | |
|---|---|
| Primary data reduction | SAXSDog, beamline data pipeline |
| Data processing | PRIMUS |
| Guinier analysis | PRIMUS |
| Inverse Fourier transformation | GNOM |
| Molecular dynamics simulation | GROMACS, CHARMM force field |
| BME weighting | Own-written python script |

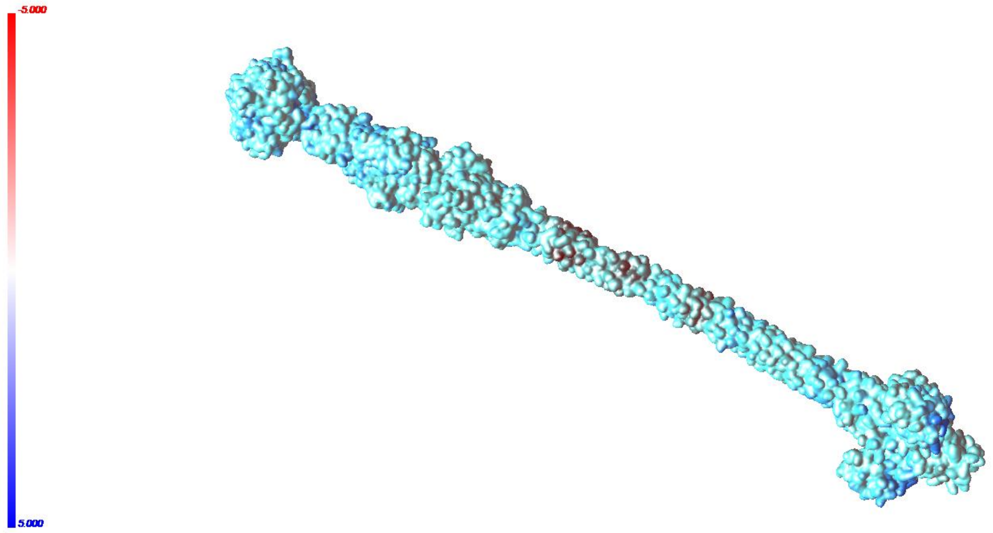


*Fig. 1. Surface charge of T7 fibers*. AlphaFold prediction of fiber surface charge, showing no differences between the two sides of the fiber. Blue color represents positive charge, red represents negatively charged surfaces.
